# Gene expression atlas of a developing tissue by single cell expression correlation analysis

**DOI:** 10.1101/477125

**Authors:** Josephine Bageritz, Philipp Willnow, Erica Valentini, Svenja Leible, Michael Boutros, Aurelio A. Teleman

## Abstract

The *Drosophila* wing disc has been a fundamental model system for the discovery of key signaling pathways and for our understanding of developmental processes. However, a complete map of gene expression in this tissue is lacking. To obtain a complete gene expression atlas in the wing disc, we employed single-cell sequencing (scRNA-seq) and developed a new method for analyzing scRNA-seq data based on gene expression correlations rather than cell mappings. This enables us to discover 824 genes with spatially restricted expression patterns, and to compute expression maps for all genes in the wing disc. This approach identifies both known and new clusters of genes with similar expression patterns and functional relevance. As proof of concept, we characterize the previously unstudied gene CG5151 and show it regulates Wnt signaling. This novel method will enable the leveraging of scRNA-seq data for generating expression atlases of undifferentiated tissues during development.

## MAIN TEXT

The *Drosophila* wing imaginal disc has been an important model system for the molecular dissection of tissue growth control, pattern formation, epithelial morphogenesis, inter-cellular communication and signaling, cell competition, and the biophysical interaction of cells^1–8^. Despite widespread use of the wing disc, the expression patterns of the vast majority of genes in the wing disc are not known. Recent advances in single-cell sequencing should enable an approach to generate genome-wide expression maps based on dissociating the wing disc, sequencing single cells, and then reassembling the wing disc *in silico* based on the expression patterns of known genes. Although single-cell sequencing has been used to identify different cell types in organ systems^9–12^, the wing disc presents several challenges. Firstly, the wing disc is composed mainly of pluripotent, undifferentiated stem-like cells, hence it consists of very few different cell types. Secondly, with 50,000 cells, the wing disc has one order of magnitude more cells than, for instance, the entire *Drosophila* embryo^13^. We present here an approach to use single-cell sequencing for generating gene expression maps in such a large and undifferentiated tissue. This approach is based on analyzing the correlation of expression between genes rather than mapping of sequenced cells back to their locations in the tissue. This allowed us to discover 824 genes with spatially restricted expression patterns in the wing, and to compute expression maps for all genes in the wing disc. This approach will enable the use of single-cell sequencing to generate gene expression maps in developing, undifferentiated tissues.

## RESULTS

To construct a gene expression map of the *Drosophila* wing disc, we dissociated cells from wing discs of 3rd instar female larvae and sequenced their mRNA using DropSeq^14^. This yielded RNA sequences from 1,468 cells with a median depth of 3,774 transcripts and 1,134 genes per cell (Supplementary Figure 1A-B), in line with or better than what others have reported for *Drosophila* cells using this method^9, 11, 13^. We could unambiguously identify true cell barcodes (Supplementary Figure 1C) indicating a low level of ambient mRNA, and hence cell breakage, during sample preparation. Gene expression values of two biological replicate DropSeq libraries correlated very highly to each other (r=0.93, Supplementary Figure 1D) indicating reproducibility of the data. The average gene expression values obtained by combining together all the single-cell reads correlated well with RNA-seq data of whole, non-dissociated wing discs (Supplementary Figure 1E), suggesting that the DropSeq data captured most of the gene expression in the wing disc and that the DropSeq procedure, including the cell dissociation, did not strongly alter gene expression in the disc.

**Fig. 1:**
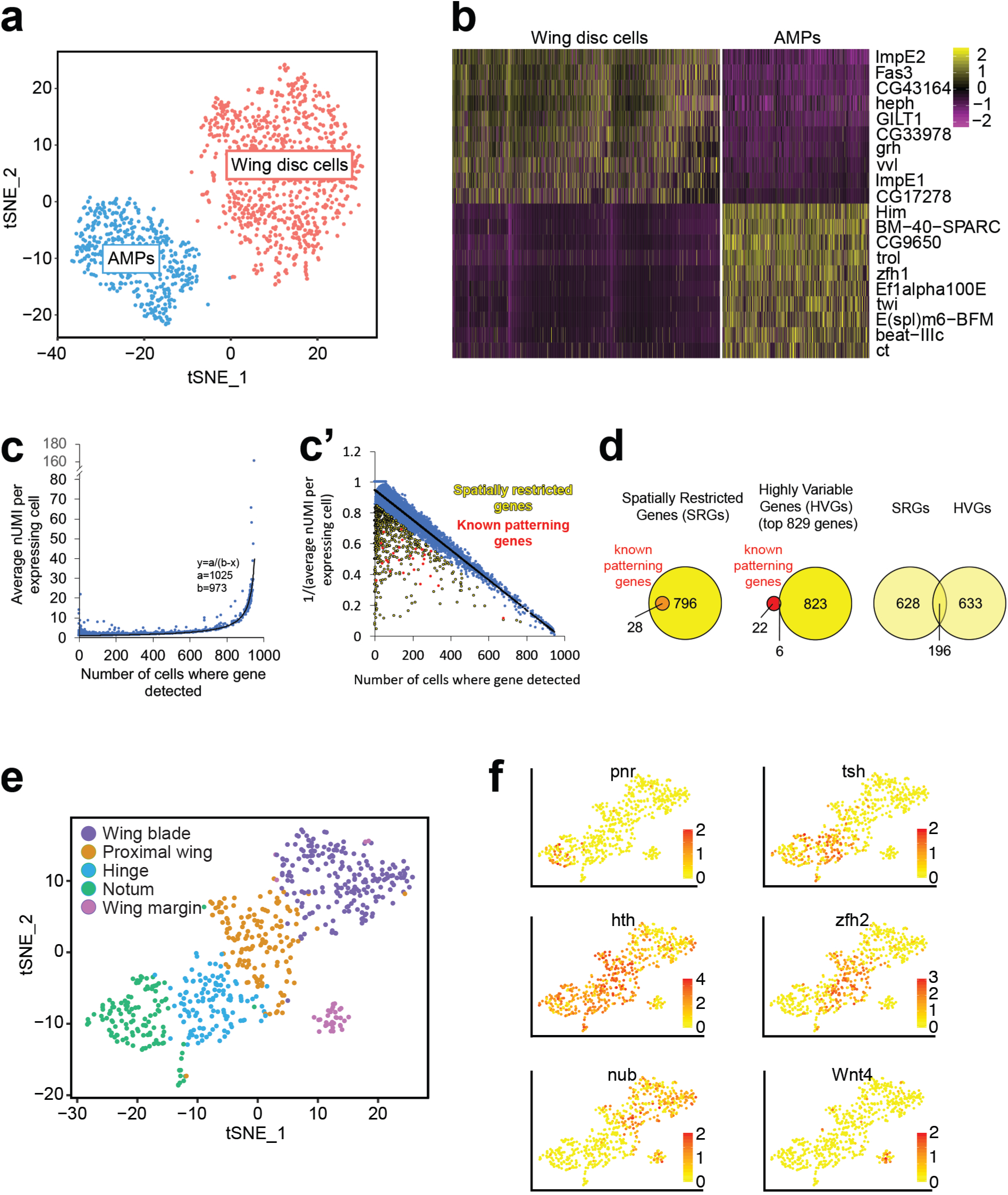
Single-cell sequencing of wing disc cells identifies Spatially Restricted Genes. **a, b**, Two-dimensional t-SNE representation of all sequenced cells reveals two main cell clusters (a) corresponding to wing disc cells and adult muscle precursors (AMPs), based on differential expression of AMP genes in the two clusters (b). **c, c’**, Identification of Spatially Restricted Genes (SRGs) as genes observed in fewer cells than expected based on their expression level. **d**, The set of 824 SRGs contains all 28 benchmark, patterning genes known to have spatially restricted expression domains based on literature (Supplementary Table 1). In comparison, a similarly sized set of ‘Highly Variable Genes’ contains only 6 of the 28. **e,f**, Two-dimensional t-SNE representation of all wing disc cells using the 824 SRGs for dimensional reduction identifies 5 clusters along the proximal-distal axis of the wing disc (e), based on expression of known marker genes (f).

To identify sub-populations of cells in the wing disc, we clustered cells using a graphical approach implemented in the Seurat R package for single-cell sequencing data^15^. Visualization by t-Distributed Stochastic Neighbor Embedding (t-SNE) identified two distinct cell populations (Fig. 1a). Inspection of the main marker genes distinguishing these populations (Fig. 1b) revealed that the two clusters correspond to cells of the wing disc proper (Fig. 1a, red) and adult muscle precursor cells (AMPs, Fig. 1a, blue) which are attached to the basal surface of the dorsal wing disc. Since we focus here on the wing disc proper, we excluded AMP cells from all subsequent analyses.

To identify genes with a spatially restricted expression patterns (which we term Spatially Restricted Genes, SRGs), we plotted for every gene the number of cells in which it was detected versus the average expression level of the gene in those expressing cells (Fig. 1c). The rationale is that for ubiquitously expressed genes, the stronger the gene is expressed, the higher the chance the mRNA will be captured by the DropSeq beads, and hence the higher the number of cells in which it will be detected. Indeed, we find that most genes lie on a curve that progressively increases and asymptotes near the total number of cells sequenced. The SRGs are the genes that are observed in fewer cells than expected, given the level of expression of the gene (i.e. the dots above the curve, Fig. 1c). These were identified as genes with residuals smaller than 1 standard deviation below the mean on the inverse graph (yellow points in Fig. 1c’), yielding a set of 824 SRGs. As a benchmark, we compiled a list of 28 genes well-known from the literature to be expressed in specific domains of the wing disc, such as engrailed, dpp, apterous, or wingless (Supplementary Table 1). The 824 SRGs included all 28 of these known patterning genes (red dots in Fig. 1c’, and Fig. 1d) and almost all genes known to have a spatially restricted expression pattern in the wing disc. In comparison, a similarly sized set of 829 “Highly Variable Genes” (HVGs) identified using the Seurat R package^15^ only contained 6 of the benchmark genes (Fig. 1d), suggesting the analysis presented here is well suited for our specific goal of identifying genes with spatially restricted expression domains.

By using this set of SRGs for dimensional reduction and clustering, wing disc cells clustered into five clusters along the proximal-distal axis of the wing, corresponding to the wing margin, the wing blade, the proximal wing, the hinge, and the notum (Fig. 1e), as could be seen by the expression levels of wing margin (Wnt4), pouch (nub), or hinge/notum genes (tsh, zfh2, pnr, hth) in the five clusters (Fig. 1f). We confirmed that there were no biases in these clusters in terms of the number of Unique Molecular Identifiers (UMIs)/cell, read alignment rate, fraction of mitochondrial RNA or representation of the two biological replicates (Supplementary Figure 2). The major wing disc regions were retrieved by our clustering approach, indicating a successful cell isolation from the entire tissue.

**Fig. 2:**
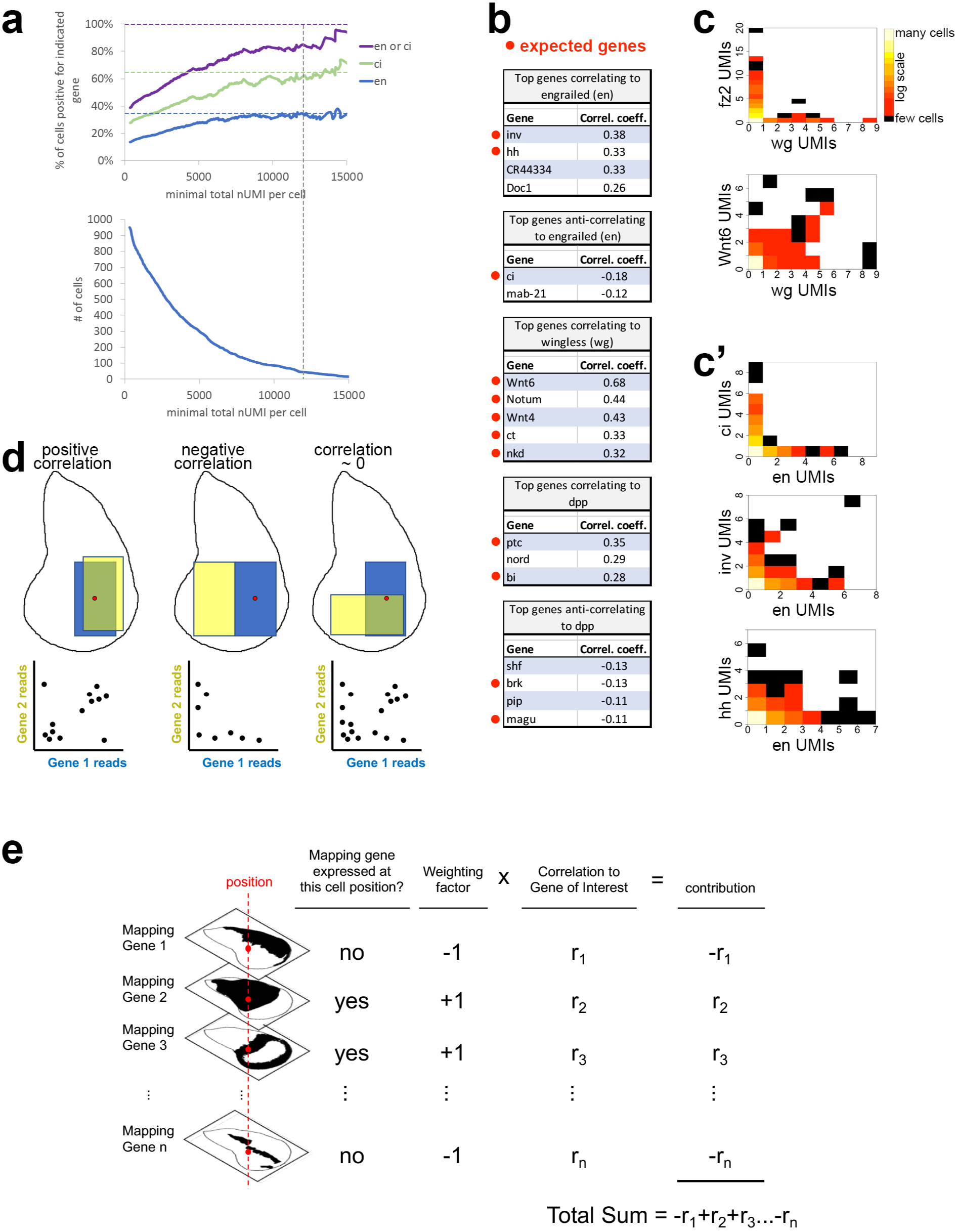
Generation of gene expression maps based on gene expression correlations. **a**, The high false-negative rate of scSeq makes it difficult to confidently conclude that a gene is not expressed in any one sequenced cell, and hence to confidently locate its original position in the wing disc. Shown is the number of cells that are positive for expression of en or ci, which have largely mutually exclusive expression patterns in the wing, at different sequencing depth thresholds. nUMI= number of unique molecular identifiers. **b**, Top hits from genome-wide correlation analysis of gene expression across all sequenced wing disc cells. **c-c’**, Two-dimensional histograms showing the distribution of all sequenced cells according to the level of expression of the two indicated genes. **d**, Concept for generating expression maps based on gene expression correlations. Positive correlation between two genes indicates they have overlapping expression domains, whereas a negative correlation indicates expression domains that are more mutually exclusive. **e**, Schematic representation of the method used to generate computed expression maps.

We first tested whether we could determine the location in the wing disc of the sequenced cells, based on the presence or absence of expression of genes with known expression patterns, such as engrailed (for the posterior of the wing), ci (for the anterior), apterous (for dorsal) and so on (Supplementary Figure 3). Since the expression pattern of many genes is known in the wing disc, the intersection of these gene expression domains could allow precise placement of sequenced cells. However, although our DropSeq data are of high quality, we found we could not confidently map the location of the sequenced cells because the transcriptome coverage of current single-cell approaches does not allow us to distinguish whether a gene is not expressed or not detected in any given cell: Although roughly 35% of wing disc cells should express engrailed (estimated by measuring the area of the engrailed expression domain of a wing disc), and 65% of disc cells should express the gene ci with complementary expression pattern, in our entire library only 14% of cells were en^+^ (>0 reads for en) and 28% were ci^+^ (left edge of graph, Fig. 2a). We tested if this could be solved by setting a minimum UMI per cell threshold. Setting a minimum requirement of 12,000 UMIs/cell, however, still resulted in only 84% of cells being en^+^ or ci^+^, and only 45 of the 948 sequenced wing disc cells passed this threshold (Fig. 2a). We therefore searched for an alternate method to leverage these data and build a wing disc expression map.

**Fig. 3:**
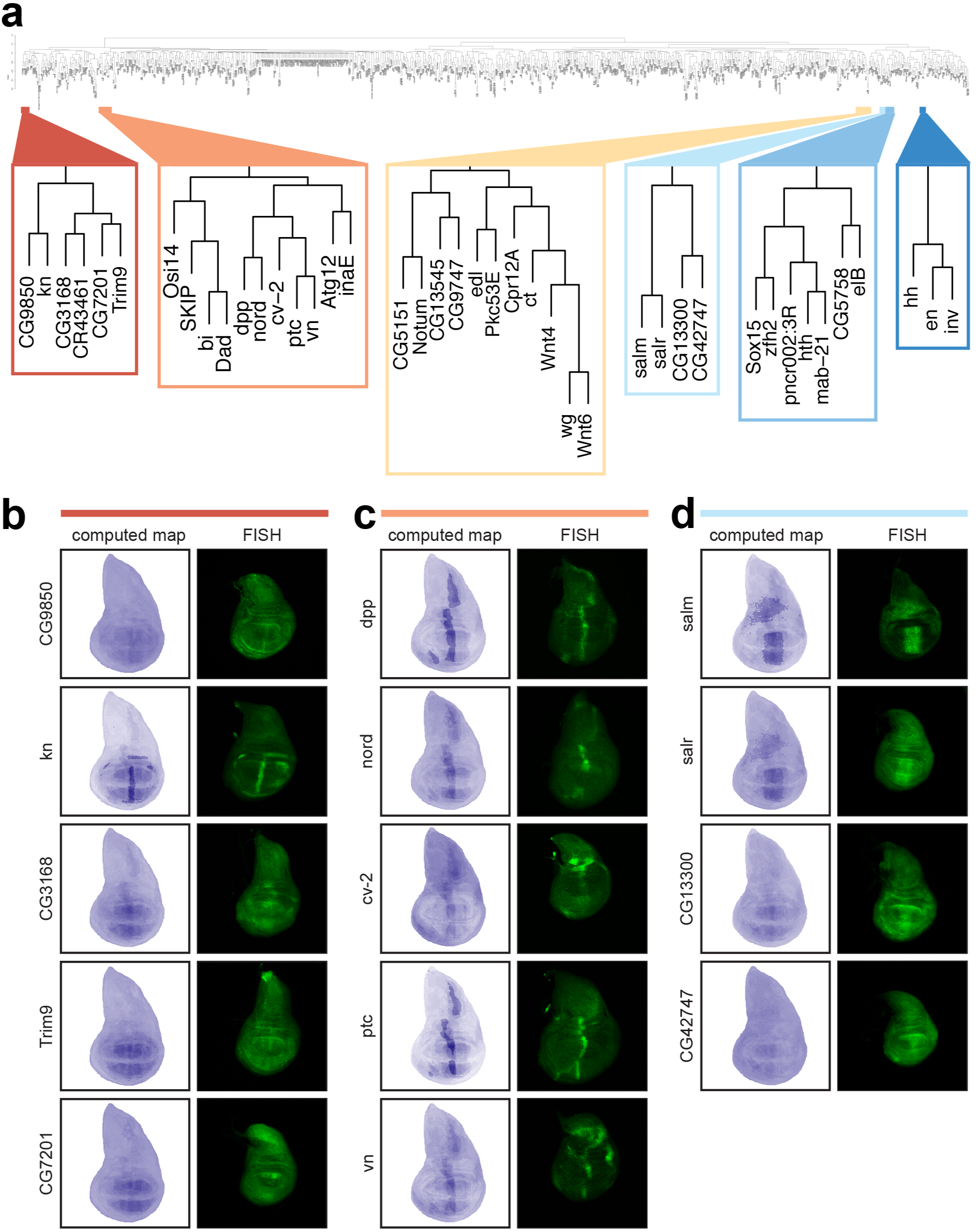
Computed expression maps for genes of unknown expression patterns agree well with their actual expression detected by *in situ* hybridization. **a**, Hierarchical clustering of SRGs by expression correlation identifies clusters of genes with related expression patterns containing both genes of known and unknown function. **b-d**, Expression patterns of genes detected by *in situ* hybridization largely confirm the expression patterns predicted by the computed maps. For testing, all genes in the ‘red’ (b), ‘orange’ (c) and ‘light blue’ clusters (d) shown in panel (a) were selected.

We noticed that correlations in gene expression between genes, based on their expression across the hundreds of sequenced cells, are quite good. For instance, we calculated correlation coefficients between en and all other genes in the genome across the sequenced cells and found that, as expected, the top genes genome-wide correlating to en are inv and hh, and the top anti-correlating gene to en is ci (Fig. 2b). Likewise, the top genes either correlating or anti-correlating to wg or dpp are also known to be expressed in either overlapping or complementary expression patterns, respectively, in the wing disc (Fig. 2b). The underlying data can be visualized using 2-dimensional histograms (Fig. 2c). For instance, in the case of wingless (wg) and frizzled 2 (fz2) which are expressed in largely complementary domains, many cells have detectable transcripts for fz2 or for wg, but few cells express both (Fig. 2c). In contrast, a good number of cells have detectable transcripts for both wg and Wnt6 (Fig. 2c), as expected given that they are expressed in overlapping domains. Likewise, few cells are en^+^/ci^+^, whereas many cells are en^+^/inv^+^ or en^+^/hh^+^, as expected from their relative expression domains (Fig. 2c’). Interestingly, this correlation analysis also identifies novel genes which correlate strongly with en and therefore likely have a similar expression pattern, such as the non-coding RNA CR44334 (Fig. 2b).

We therefore conceived a method for generating gene expression maps based on gene correlations, which does not necessitate mapping the location of the sequenced cells in the tissue. This method uses the concept that the correlation coefficient between two genes indicates whether the expression domain of the two genes is overlapping (positive correlation), complementary (negative correlation), or orthogonal (no correlation) (Fig. 2d). Therefore, for a given cell within the expression domain of Gene 1 with known expression pattern (red dot, Fig. 2d), uncharacterized Gene 2 is likely also expressed if the two genes correlate, and not expressed if they anti-correlate. If the correlation coefficient is close to zero, then the expression domain of Gene 1 is not informative with regards to Gene 2. We therefore compiled a virtual map of the wing disc containing the expression domains of 58 genes known from the literature to have distinct expression patterns which we term ‘mapping genes’ (Fig. 2e), and we calculated a cross-correlation matrix between these 58 mapping genes and all genes in the genome. To compute an expression map of a gene, for each position in the wing disc we then added the correlation coefficients between this gene and the mapping genes with a weighting factor of either +1 or −1 depending on whether the mapping gene is expressed in that position or not (Fig. 2e). We call these maps ‘computed expression maps’. We then tested this approach by performing fluorescent *in situ* hybridizations (FISH) to assay whether the computed maps and the *in situs* agree with each other (Fig. 3-4).

The method described above generates computed expression maps for all genes in the genome. Based on these, there are multiple different ways to sort out genes of interest based on similarity of their expression patterns to known genes of interest. Here we present three different ways: 1) clustering genes using a two-dimensional dendrogram, 2) searching for genes that correlate or anti-correlate with one specific gene of interest, and 3) generating an interaction network based on gene expression similarities. To cluster genes by expression pattern, we calculated a cross-correlation matrix of gene expression for all 824 SRGs against each other, and then used this to hierarchically cluster the genes according to their expression patterns (Fig. 3a and Supplementary Data 1 for a high-resolution dendrogram). Visual inspection of this dendrogram confirmed that genes that cluster together have similar expression patterns in the wing. For instance, the ‘dark blue’ cluster (Fig. 3a) consists of en, inv and hh, which are co-expressed throughout the posterior compartment of the wing disc. The ‘medium blue’ cluster consists of genes expressed in the proximal region of the wing disc such as hth and zfh2, together with other genes of unknown expression pattern or function. The ‘yellow’ cluster consists of wg, Wnt4, Wnt6 and cut (ct), which are all expressed on or near the dorsal/ventral boundary of the wing disc. We selected three clusters which contain both characterized and uncharacterized genes and performed *in situs* on all genes in the cluster. The ‘red’ cluster (Fig. 3b) consists of genes enriched in the wing pouch with a pattern along the anterior-posterior axis. The gene ‘kn’ is one of the 58 ‘mapping genes’ hence the computed map matches the *in situ* because it is one of the inputs into the mapping algorithm. CG9850, a gene of unknown function and expression pattern, is predicted by the computed map to also have a mild ‘kn-like’ stripe that is less accentuated than kn, and indeed this matched the fluorescent *in situ*. The uncharacterized gene CG3168 was predicted according to the computed map to have a broader expression pattern in the wing pouch that is repressed at the dorsal/ventral boundary (Supplementary Figure 3). Indeed, this pattern was confirmed by *in situ* hybridization (Fig. 3b). The gene Trim9, involved in neurogenesis in the central nervous system ^16^, but of unknown expression pattern or function in the wing, was predicted to be expressed predominantly in the wing pouch with an inverse venation pattern and inhibition at the D/V boundary. This complex expression pattern was also confirmed by *in situ* hybridization (Fig. 3b). The *in situ* for the last gene in the cluster, CG7201, had some elements of the predicted map, such as higher expression medially and broad repression at the D/V boundary, but it also differed somewhat from the map. Thus, overall, the computed maps are able to predict the main features of the gene expression patterns. Along the same lines, we performed *in situs* for genes in the orange and light-blue clusters, and the *in situs* confirmed the broad characteristics of the computed maps (Fig. 3c, d). Interestingly, due to their expression patterns, this implicates a number of genes with previously uncharacterized functions and/or expression patterns in anterior-posterior patterning and ptc or dpp signaling. For instance, the expression pattern of the functionally uncharacterized gene nord is largely overlapping with the expression patterns of dpp or ptc (Fig. 3c).

A second way to identify genes of interest is to select genes that have expression patterns that either correlate or anti-correlate with a specific gene of interest, such as senseless (Supplementary Table 2), wg (Supplementary Table 3) or Dpp (Supplementary Table 4). Amongst these are many genes that have previously been implicated in the respective signaling pathways (Supplementary Table 2-4 and Fig. 2b). Hence, we only performed *in situs* for the top correlating/anti-correlating genes with distinct expression patterns that had not previously been characterized in the wing disc. Fig. 4a shows *in situs* for genes that correlate with the neurogenic gene senseless, which is expressed along the dorsal/ventral boundary of the wing and in clusters of sensory precursor cells in the body wall (arrowheads). In all cases, the *in situ* confirmed the pattern predicted by the computed maps (Fig. 4a), thereby implicating novel genes in wing neurogenesis. For instance, Fhos is involved in actin stress fiber formation ^17^ and hence may play a role in neurogenesis, and ImpL3 is the metabolic enzyme lactate dehydrogenase. Rau and cpo have previously been implicated in neurogenesis in other organs ^18–20^. Amongst the genes correlating with wingless (Fig. 4b), CG10249 (Kank) has been linked to attachment sites between muscle and epidermal cells in the embryo ^21^. One additional uncharacterized gene correlating with Dpp is CG9689 (Supplementary Figure 4).

**Fig. 4:**
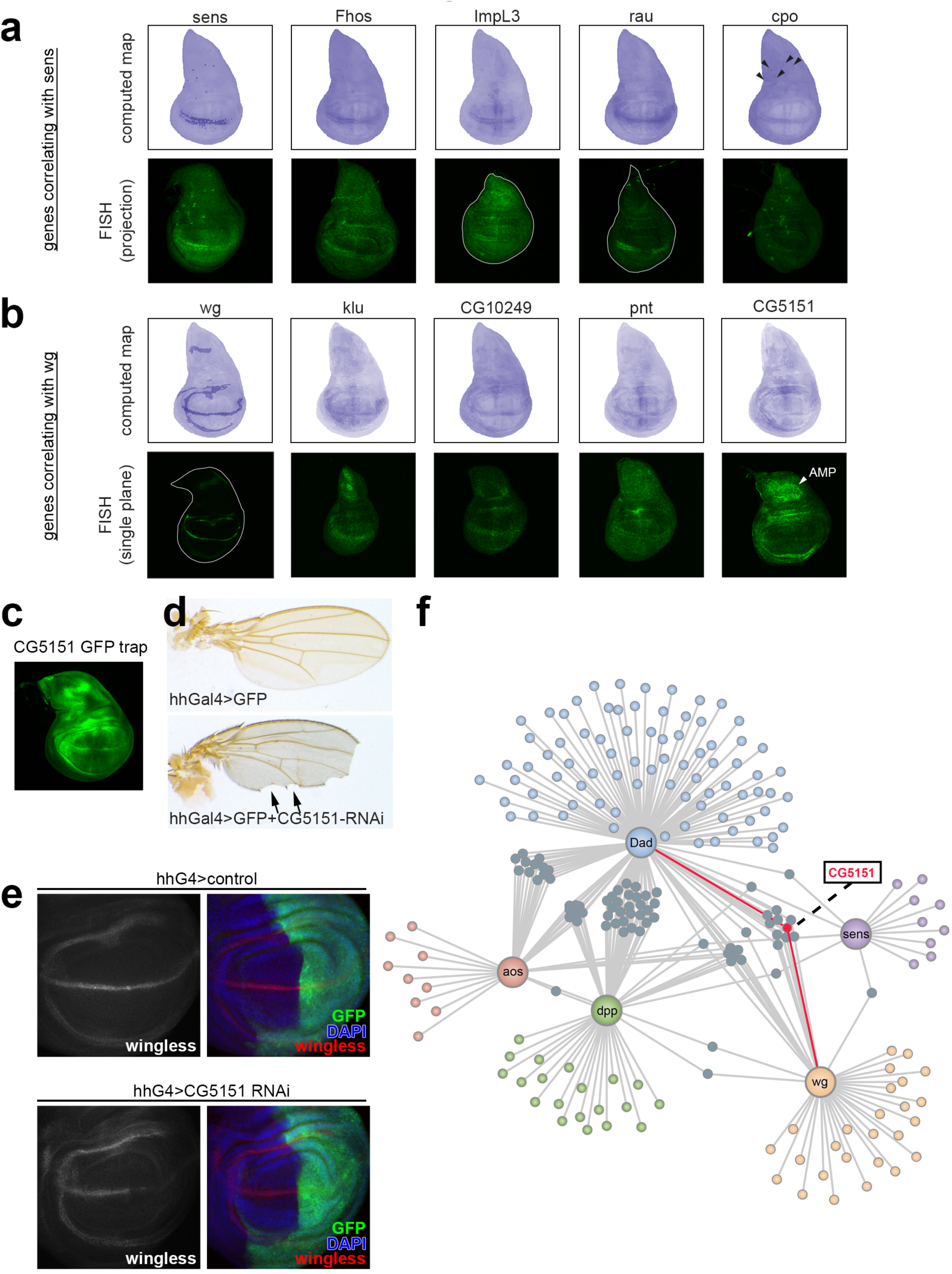
Discovery of genes in processes of interest based on their expression pattern. **a, b**, Computed expression maps and *in situ* hybridizations for genes correlating with either senseless (a) or wingless (b). **c**, Expression of CG5151 using a GFP transcript trap in the endogenous CG5151 locus reveals expression at the D/V boundary and in a more proximal ring, similar to that of wingless. **d, e**, Knockdown of CG5151 in the posterior compartment of the wing disc using hedgehog-Gal4 (hhGal4) causes notching of the posterior wing margin, a typical notch or wingless loss-of-function phenotype (d) and loss of wingless protein (e). **f**, Gene-network analysis of the most connected genes linked to Dpp, Wnt, Notch and/or EGRF signaling pathways. Only edges with a minimum correlation coefficient of 0.1 are shown. Self-correlations are excluded.

We selected CG5151 as a gene to study in more detail, as it is functionally uncharacterized and has a human ortholog LDLRAD4 (also known as C18ORF1). The computed map predicts CG5151 to be expressed weakly along the dorsal/ventral boundary and in a more proximal ring, coinciding with wingless expression. *In situ* hybridizations confirmed this expression pattern, and also detected expression of CG5151 in the Adult Muscle Precursor cells, AMPs, which are not part of the disc proper and are not included in our computed maps (Fig. 4b). This expression pattern was also observed with a GFP transcript trap in the endogenous CG5151 locus (Fig. 4c). We next tested if CG5151 might be involved in wingless or notch signaling. Knockdown of CG5151 in the posterior half of the wing caused wing notching (a phenotype typical for Notch or wingless loss-of-function, arrows in Fig. 4d), and strongly reduced wingless expression (green domain in Fig. 4e). In sum, as proof of principle, our mapping strategy allowed us to identify a novel uncharacterized gene, CG5151, which has an expression pattern that overlaps with that of wingless, and is functionally involved in wingless/notch signaling. Interestingly, the human ortholog LDLRAD4 is functionally not well characterized but its expression is elevated in hepatic cancers and it promotes tumorigenesis ^22^. It will be interesting to test whether Wnt or Notch signaling are involved in its tumorigenic activity.

## DISCUSSION

The quality and resolution of the expression maps depends on several parameters which can be further refined. One parameter is the alignment of the ‘mapping gene’ maps to each other (left side, Fig. 2e). This is non-trivial because each ‘mapping gene’ map derives from an *in situ* on an individual wing disc, and no two wing discs have exactly the same morphology. Furthermore, every map needs to be aligned to every other map, which is a problem that scales exponentially with the number of mapping genes. Secondly, the quality of the computed maps will increase with the number of single cells sequenced (which influences the expression cross-correlations) and with the number of mapping genes. That said, our current setup with 58 mapping genes and 948 cells sequenced is sufficient to yield good quality computed maps that agreed well with the *in situ* hybridizations we have done.

In sum, we identify here 824 Spatially Restricted Genes in the wing disc, thereby in a single sc-Seq experiment increasing the number of known genes with restricted expression pattern in the wing by roughly ten-fold. We developed a method based on gene expression correlation analysis to obtain computed expression maps for these 824 Spatially Restricted Genes (Supplementary Data 2) which agree well with *in situs* for the genes we have tested. This work also yields a network of genes with expression patterns linked to signaling pathways of interest such as Dpp, Wnt, Notch or EGRF (Fig. 4f and Supplementary Figure 5 with gene names) which identifies novel genes such as CG5151 that are functionally linked to these pathways. This approach can be applied more broadly to generate expression maps of genes in developing undifferentiated tissues.

## MATERIALS & METHODS

### Drosophila stocks

The following fly lines were used: w^1118^, CG5151^RNAi^ (VDRC ID 102217), CG5151 MiMIC (Bloomington stock 52188). Stocks were maintained at 25 °C with a 12 h light/dark cycle, except for the crosses used in the knockdown experiments with RNAi and GAL4/UAS expression, for which crosses were maintained at 29 °C.

### Single cell sample preparation from wing disc tissue

Wing discs of female wandering 3rd instar w^1118^ larvae were dissected in Schneider’s medium in batches of 5 animals (to prevent hypoxia) and transferred into a tube containing Schneider’s medium on ice for a maximum time of 30 minutes. The isolated wing discs were rinsed once with Schneider’s medium and then incubated for 15 minutes in a water bath at 37°C in TrypLE (10x), with gentle mixing every 5 minutes. Schneider’s medium was then added to the loosened tissue pellets, followed by gentle mechanical dissociation using a P1000 pipette. The cell suspension was then passed through a 10 μM cell strainer to remove undigested tissue and cell clumps. Cells were manually counted using a plastic hemocytometer (C-Chip N01). The entire cell isolation protocol was done with PBS-Triton (0.1%) coated microcentrifuge tubes and tips to minimize cell loss.

### scRNA-seq by Drop-Seq technology

Drop-Seq experiments were performed as published ^14^ following the detailed online protocol (Drop-seq-Protocol-v1.0-May-2015). In brief, cells and barcoded beads (ChemeGene) were co-flown in an Aquapel coated microfluidics device (FlowJem) and co-encapsulated in aqueous droplets for a maximum period of 15 minutes. Isolated wing disc cells were loaded without further dilution at a concentration found by species mixing experiments to contain a maximum of 3% cell doublets. The aqueous flow rates were adjusted to ensure stable production of monodispers droplets. The size of the droplets was controlled by the oil flow rate. For this project, settings were chosen to generate about 120 μM droplets for batch 1 and 85 μM droplets for batch 2. While the standard barcoded beads were used for batch 1, batch 2 was performed with filtered beads (< 40 μm in diameter) to account for the smaller droplet size. High quality emulsions were broken by perfluorooctanol and reverse transcription of captured mRNA was started immediately after. Subsequently, barcoded beads were incubated with Exonuclease I to remove excess primers, and cDNA was then amplified from 2000 beads per reaction (12-14 PCR cycles). Up to 10 reactions were pooled, purified with a 0.6 ratio of AMPure beads (Agencourt) and eluted in the necessary amount of water to obtain 400-1,000pg/ul of cDNA. Final libraries were prepared using the Illumina Nextera XT kit and 1 ng of amplified cDNA as input. The average size of sequenced libraries was between 700 and 800 bp. Paired-end sequencing was carried out with the Illumina HiSeq2500 instruments at the DKFZ Genomics and Proteomics Core Facility (Heidelberg, Germany).

### Preprocessing of Drop-Seq data

Paired-end sequence reads were processed as described ^14^. The available R command lines were implemented in our in-house Galaxy^1^ server (http://galaxy-b110.dkfz.de/galaxy/) following the default settings described in detail in the Drop-seq computational cookbook (http://mccarrolllab.com/wp-content/uploads/2016/03/Drop-seqAlignmentCookbookv1.2Jan2016.pdf). The reads were aligned to the *Drosophila* reference genome (BDGP6 version 87 (GCA 000001215.4)) using STAR 2.5.2b-0 with the default parameters. The cell number was estimated by plotting the cumulative fraction of reads per cell against the sorted cell barcodes (decreasing number of reads) and determining the point of inflection. The raw digital gene expression matrices were generated for the two batches. Further filtering of the expression matrices was done to ensure high-quality single-cell data. By using the Seurat R package^2^, we selected cells with low expression of mitochondrial encoded genes (<5%), high alignment rate (>85%) and a minimum number of detected genes (>200). Outlier cells (>3,000 detected UMIs), which could be potential cell doublets, were also excluded from further analysis. After subsetting the wing disc cell population, we also applied a reasonable UMI cutoff (>2,000). The UMI cutoff was empirically determined by performing correlation analysis with genes of known expression patterns. Using cells with at least 2,000 detected UMIs showed the expected correlation coefficient values among our reference set. Additionally, we also removed genes that were detected in only 1 cell. This filtering resulted in 615 high-quality wing disc single cells, which were subsequently merged together in a single DGE matrix. Prior to Principle component analysis and clustering, the data were log transformed (log +1) and re-scaled by multiplying by 10,000.

### Identification of spatially-restricted genes

Spatially Restricted Genes (SRGs) were identified by analyzing the nUMIs, as this led the smallest spread in the data. A scatter plot was generated for all detected genes whereby the x-axis is the number of cells in which the gene was detected (nUMI > 0) and the y-axis is the average nUMI for that gene in the cells in which it was detected (i.e. not across the entire cell population, since this also contains cells not expressing the gene). A linear model was then fitted to the data, and residuals were calculated for each gene relative to the linear model. The average and standard deviation of the residuals was calculated, and SRGs were defined as genes with residuals < (mean – 1 std dev).

### Batch Correction, Principle component analysis and clustering

For cell/cluster identification we applied the Seurat R package ^15^ and followed largely the tutorial instructions from the Seurat website http://satijalab.org/seurat/. In order to reduce dimensionality, principle components analysis (PCA) was run on the entire transcriptome after scaling and centering the data and removing technical confounder factors (number of UMIs, number of genes and alignment rate). We examined the effectiveness of removing confounder effects by *i.)* inspecting the t-SNE plots for evenly distributed batches/libraries, number of genes and transcripts, and fraction of mitochondrial RNA among the clusters (Supplementary Figure 2D), *ii.)* analyzing the loading of genes (“PC loading”) of the different batches/libraries for their similarity and *iii.)* comparing the inter-batch correlations (Supplementary Figure 1D). The jack straw statistical analysis [num.pc = 20, num.replicate = 1,000, prop.freq = 0.01] and plotting the eigenvalues in decreasing orders (‘Elbow plot’), was used to select PCs as input for clustering. We used t-SNE for visual representation of the clusters and highlighting marker gene expression. Two distinct clusters of cells were identified by means of the above described procedure, one of which was strongly defined by expression of adult muscle precursor (AMP) marker genes. As this cell type was irrelevant for the present study, we excluded it based on the t-SNE plot from further analysis. After AMPs were removed, the list of spatially-restricted genes (SRGs) derived specifically for the wing disc cell population was used for dimensional reduction. Selection of principle components and clustering was again performed as described above. The proportion of wing disc cells within each cluster was found to be similarly represented in both batches underlying the robustness of the identified clusters.

### Generating bulk mRNA-seq data

Wing discs from 50 female wandering 3rd instar w^1118^ larvae were dissected in Schneider’s medium, 5 larvae at a time to prevent hypoxia, and transferred to a tube containing Schneider’s medium on ice. The wing discs were then lysed in TRIzol (Thermo Fischer) for total RNA isolation following manufacturer’s protocol. RNA library preparation (TruSeq Stranded mRNA Sample Preparation Kit, Illumina) and sequencing (50 nt single-end reads, HiSeq2500, Illumina) were done at the DKFZ Genomics and Proteomics Core Facility (Heidelberg, Germany) following the manufactures’ protocol. Fastq sequencing data was processed using the scater R package ^23^ by applying the default settings.

### Comparison of single-cell and bulk transcriptomic profiles

To compare gene expression data at single-cell and bulk levels, we calculated Pearson’s correlation coefficients (R) of gene expression for all possible gene pairs across the cells. Only genes detected in both sets were used for comparison. For single-cell data, the average UMI expression for each gene was first calculated and then converted to average transcripts per million (ATPM). Gene counts were converted to TPM (Transcripts per million) and isoform counts averaged. Log-transformed data was plotted (1 + ATPM/TPM).

### Calculation of gene expression correlations

For gene expression correlation analysis, only cells with nUMI>2,000 were considered, since cells with fewer reads per cell reduced the correlation coefficients. Gene expression correlations across cells were calculated using the Pearson’s correlation coefficient, except that the one single cell contributing most strongly to the correlation coefficient was removed to avoid outliers from influencing the correlation. Specifically, for genes x and y, where the nUMI for each gene in cell i are x_i_ and y_i_ respectively, the means 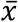 and 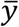 across cells were calculated. Then for every cell, the numerator of the Pearson’s coefficient 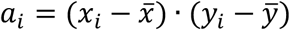 was calculated. The cell with the maximum *a*_*i*_ was excluded, and the Pearson’s correlation coefficient was calculated for all other cells.

### Generation of gene clustering dendrogram

The gene clustering dendrogram was generated by first computing a cross-correlation table for all SRGs using the correlation function described in the previous section, and then clustering and plotting using the R hclust() function.

### Generation of computed wing disc maps

To generate computed gene expression maps, we first performed *in situ* hybridizations of reference genes with published expression patterns, which we term ‘mapping genes’. Genes for which we could not confirm the expression pattern were discarded, yielding a list of 58 confirmed mapping genes (for list see Supplementary Table 5). For each mapping gene, we then selected one representative image, either from our *in situs* or from the published literature, depending on which had better signal. These images were morphed in Photoshop using the “Puppet Warp” function to fit one reference wing disc shape, and then the images for all 58 mapping genes were aligned to each other. Images were then thresholded using ImageJ to obtain binary images, thereby defining an expression domain for each mapping gene. Computed expression maps were then calculated as follows. We call {m_1_, m_2_, …, m_58_} the 58 mapping genes, x the gene of unknown expression pattern for which the expression map is being computed, and {c_1_, c_2_, …, c_50,000_} the 50,000 cells in the wing disc. Following the calculation described in the section above, a modified version of the Pearson’s correlation which excludes one outlier was calculated for gene x relative to each of the 58 mapping genes, yielding 58 correlation coefficients {*r*_*x*,1_, *r*_*x*,2_, … *r*_*x*,58_}. Computation of the expression map consists of determining an expression level e for gene x in each cell c_i_:

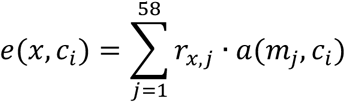

where the parameter a(m_j_,c_i_) is equal to +1 if the mapping gene m_j_ is expressed in cell c_i_, and it equals −1 if it is not. This essentially sums together all the correlation coefficients of gene x relative to the 58 mapping genes with a weighting factor of ±1 depending on whether that mapping gene is expressed in that cell or not.

### Immunostainings

Immunostainings of wandering 3rd instar wing discs were performed as previously described^3^, using monoclonal mouse anti-Wingless (clone 4D4, 1:50, Developmental Studies Hybridoma Bank) and guinea pig anti-Senseless antibody (1:300, ^24^). Secondary antibody staining was performed using fluorescently labeled antibodies at a dilution of 1:500, together with Hoechst 33342 (1:2,000, Invitrogen™) nucleic acid staining. The specimens were mounted in Vectashield mounting medium (Vector Laboratories) and imaged with a Leica TCS SP8 confocal microscope (Leica). Images were analyzed and processed in ImageJ.

### Fluorescent in situ hybridization

*In situ* probes were designed to encompass a length of 250 to 500 nucleotides and to detect all transcript variants of the gene of interest. A DNA probe template containing a T7 promoter sequence (included in the reverse primer oligonucleotide) was generated by PCR from cDNA and used to generate digoxigenin-labeled RNA probes by *in vitro* transcription reaction using the DIG RNA labeling Kit (Roche). The DNA template was removed by DNase I digest and the *in situ* probe purified from the reaction solution (RNA Clean-up, Macherey-Nagel). The purified *in situ* probe was stored in 50% formamide at − 20°C until further use.

For fluorescent *in situ* hybridization, wandering 3rd instar w^1118^ larvae were dissected and fixed in 4% paraformaldehyde for 30 min. Fixed larvae were washed 3 times in phosphate-buffered saline (PBS) containing 0.1% Tween 20 (PBT) for 10 min before dehydration in methanol/PBT (1:1) for 5 min and rinsing in methanol. Next, the larvae were incubated in methanol/PBT (1:1) for 5 min and fixed again in 4% paraformaldehyde containing 0.1% Tween 20 for 20 min. Following three washing steps with PBT for 5 min each, the sample buffer was changed to hybridization solution (HS) (50% formamide, 5x SSC (0.75 M sodium chloride and 75 mM sodium citrate dehydrate), 50 µg/ml heparin, and 0.1% Tween 20) by serial washing steps of 5 min each in HS/PBT dilutions of 30/70, 50/50, and 70/30 (vol/vol). Before hybridization, the larvae were washed for 5 min and 10 min in HS, followed by blocking for 2 h at 65°C in HS supplemented with 100 µg/ml salmon sperm DNA (AppliChem). Hybridization was performed overnight at 65°C with an *in situ* probe concentration of 1.5 ng/ml. The probe was denatured at 80°C for 5 min and cooled on ice prior to adding to the tissues. The following day, the larvae were washed with HS for 5 min and 15 min before changing the washing buffer to PBT through serial washing steps of 5 min each in HS/PBT dilutions of 70/30, 50/50, and 30/70 (vol/vol). Next, the larvae were rinsed and washed three times with PBT for 15 min and blocked in either PBT containing 5% (w/vol) bovine serum albumin or in maleic acid buffer (1M maleic acid, 1.5M NaCl; pH 7.5) supplemented with 0.5% (w/vol) blocking reagent for nucleic acid hybridization and detection (Roche) for 30 min. Binding of the antibody to the *in situ* probe was performed overnight at 4°C in the respective blocking solution containing pre-absorbed anti-digoxigenin Fab fragments conjugated to horseradish peroxidase (Roche) (1:1,000). Unbound Fab fragments were removed by rinsing three times with PBT and washing with PBT for 10 min before staining cell nuclei with DAPI (1:2,000 in PBT) for 15 min. After removal of residual DAPI by washing with PBT for 10 min, localization of the *in situ* probe was visualized using the TSA Plus Fluorescein Kit (PerkinElmer) according to the manufacturer’s protocol. For this, the larvae were incubated with the TSA working solution for 3 min or 7 min when blocked with bovine serum albumin or with blocking reagent for nucleic acid hybridization and detection, respectively. Lastly, larvae were rinsed and washed twice with PBT for 10 min. Wing imaginal discs were mounted and analyzed by confocal microscopy. Projections of wing discs shown in this study were generated using the ‘sum projection’ function in ImageJ.

### Network analysis

For constructing a network and detecting modules, the cross-correlation matrix between core signaling pathway components in the wing imaginal disc (Dad, sens, aos, dpp, wg) and the set of SRGs was calculated. A correlation coefficient cutoff of >0.1 was applied to the network, self-correlations were excluded. Visualization of the network was done using the Cytoscape software v3.6.1 ^25^.

### Data Availability

All data generated or analysed during this study are included in this published article (and its supplementary information files).

### Code Availability

All custom code is provided in the Supplemental Information file.

## Supporting information

## ACKNOWLEDGEMENTS

We thank the High Throughput Sequencing group of the DKFZ Genomics & Proteomics Core Facility for providing excellent next-generation sequencing services. We thank Eugen Rempel for implementing the DropSeq computational cookbook on our Galaxy Server. J.B. was supported by a research stipend from the Fritz Thyssen Foundation. Research in the labs of M.B. and A.A.T. is supported by ERC Grants of the European Commission.

## AUTHOR CONTRIBUTIONS

JB, PW, EV, SL and AAT performed experiments. JB, PW, EV, MB and AAT analyzed data. JB, PW, MB and AAT wrote the manuscript.

