## Supplementary material for "Gene expression atlas of a developing tissue by single cell expression correlation analysis"

Bageritz et al.

#### Legends to Supplementary Figures

**Supplementary Figure 1: Quality and reproducibility of the Drop-Seq data**

**a, b** Violin plots showing the frequency and range of gene numbers (a) and transcript numbers (b) detected per cell for the entire set of sequenced cells.

**c,** Identification of ‘true’ cells in a pool of amplified beads based on the cell barcodes ordered by their number of reads (decreasing), and the plotted cumulative fraction of reads. The figure shows a representative knee plot for one batch.

**d, e,**Correlation of averaged gene expression between independent Drop-Seq experiments (d) and bulk RNA-seq data (e). The Pearson correlations are shown.

**Supplementary Figure 2: Uniformity of scSeq parameters across wing disc clusters**

**a, d,** Assessment of wing disc cluster quality by plotting the number of UMIs (a), the Batch ID (b), the alignment rate (c), and the fraction of mitochondrially encoded RNA (d) on the clusters identified in Fig. 1e.

**Supplementary Figure 3: *Drosophila* wing disc axes, compartments and selected gene expression patterns**

**a,** Diagrams illustrating the principle axes of the *Drosophila* wing disc (anterior-posterior, dorsal-ventral, and proximal-distal).

**b,** Diagrams showing the expression domains of several well-known patterning genes.

**Supplementary Figure 4: Identification of an uncharacterized gene with expression pattern correlating to that of Dpp.**

**a,** Computed expression maps and *in situ* hybridizations for dpp and the uncharacterized correlating gene CG9689.

**Supplementary Figure 5: Higher resolution version of the interaction map shown in Main Fig. 4f, containing gene names.**

### Supplementary Figure 1

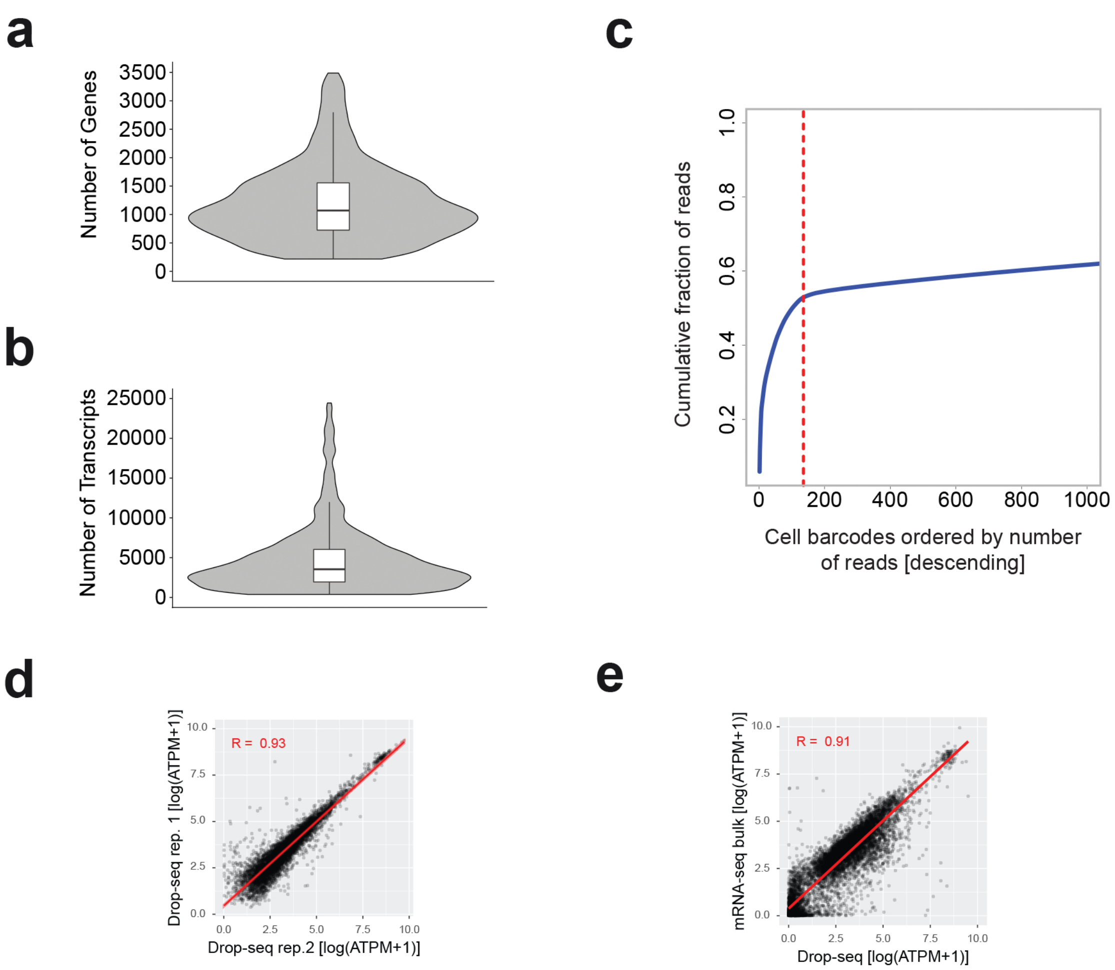

### Supplementary Figure 2

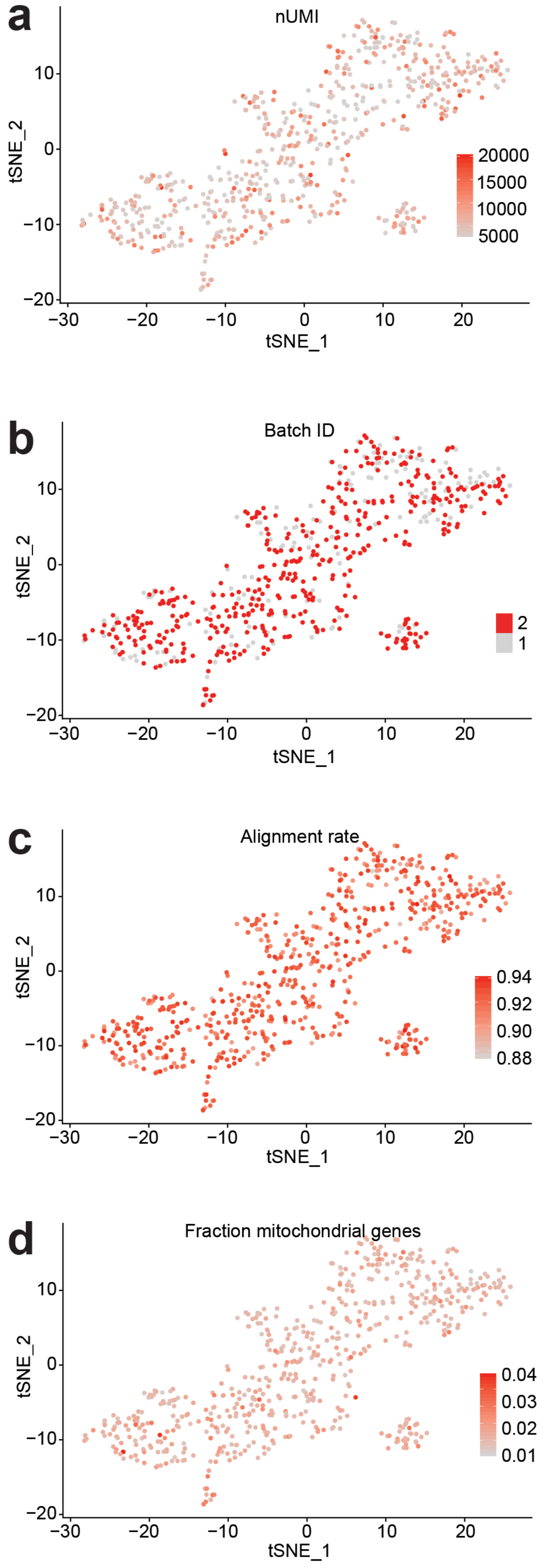

Supplementary Figure 3

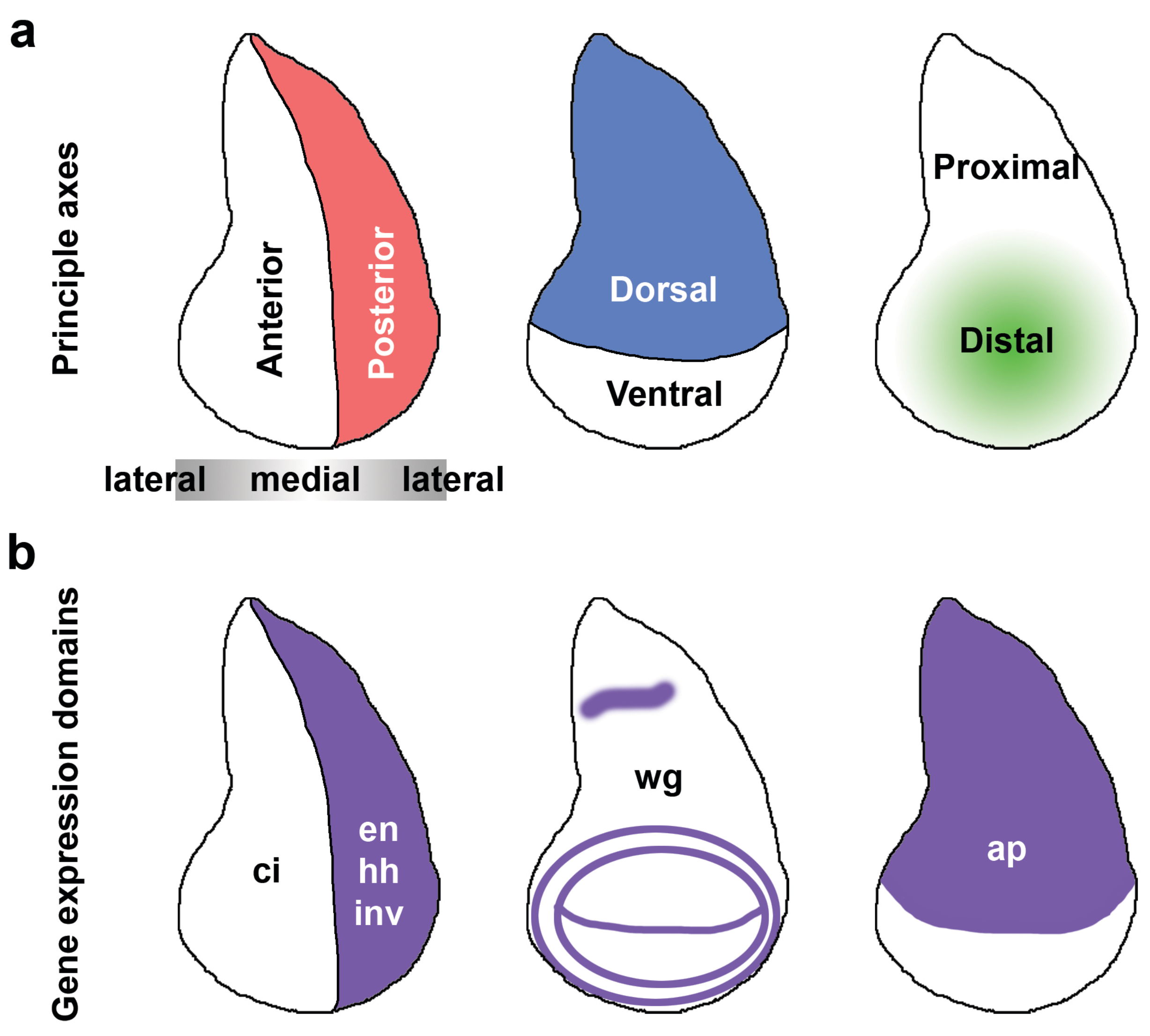

### Supplementary Figure 4

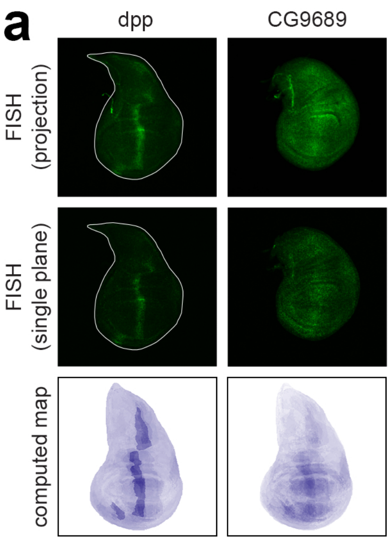

### Supplementary Figure 5

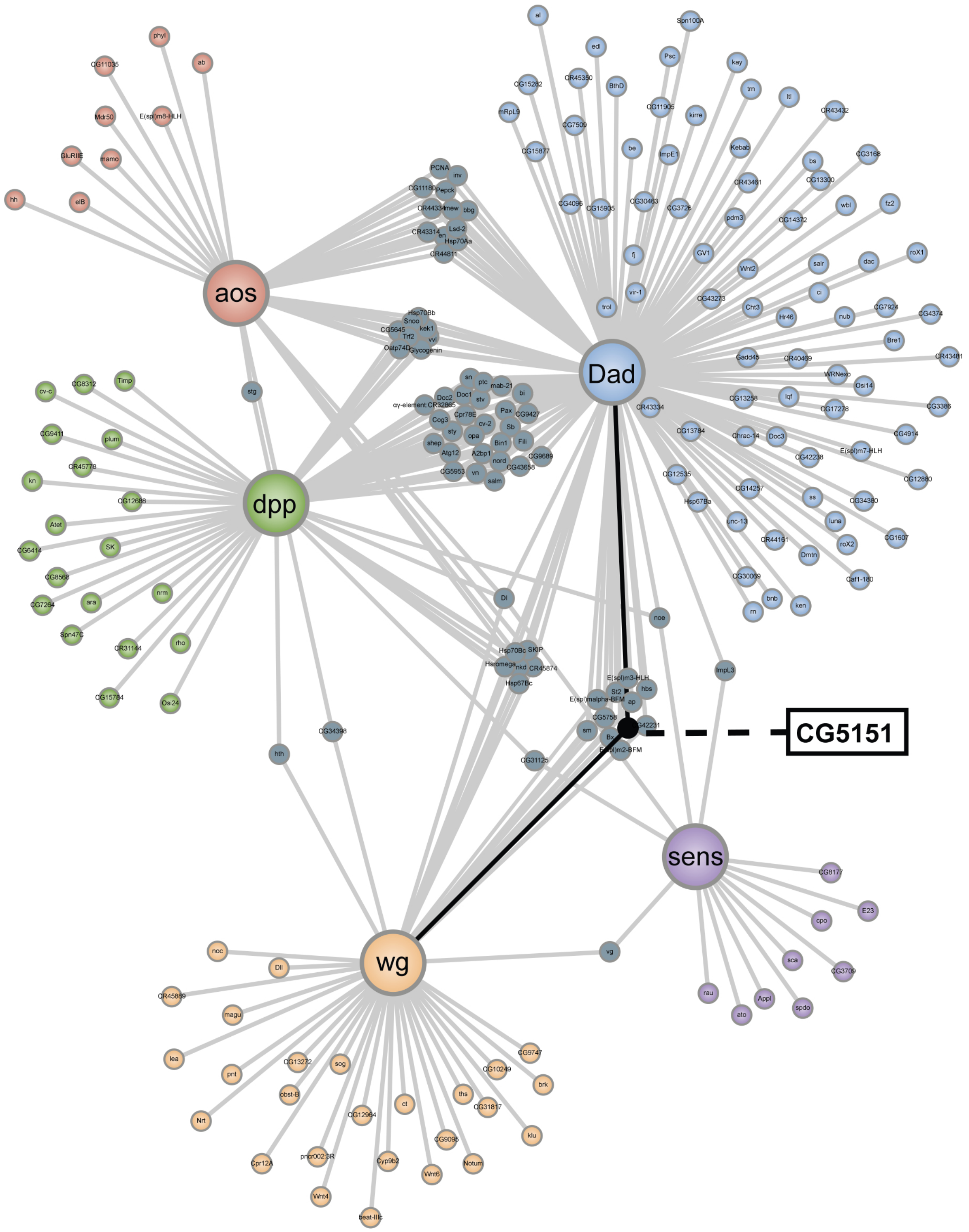

#### Custom Code

##### Code to calculate correlations between all genes and mapping genes:

```
#include <stdio.h>
#include <stdlib.h>
#include <string.h>
#include <unistd.h>
#include <math.h>

int main(int argc, char* argv[]){
	FILE *in, *in_mapping_genes, *out, *out_genes;
	int cell_num;
	int gene_num;
	int mapping_genes_num;
	int i,j,k,l;
	int found;
	char c;
	char *gene_names;
	float *reads;
	float correl;
	float mean_i, mean_j, numerator, denom_i, denom_j;
	int outlier;
	float max;

	struct correspondance {
		char name[40];
		int num;
		int num_in_seq_database;
	} *mapping_genes;

	in=fopen("expression_data.txt","r");
	if(in==NULL){
		printf("[ERROR] Could not open input file
expression_data.txt. Terminating.\n");
		exit(1);
	}

	in_mapping_genes=fopen("mapping_genes.txt","r");
	if(in_mapping_genes==NULL){
		printf("[ERROR] Could not open input file
mapping_genes.txt. Terminating.\n");
		exit(1);
	}

	out=fopen("out_cross_correlation.txt","w");
	if(out==NULL){
		printf("[ERROR] Could not open output file
out_cross_correlation.txt. Terminating.\n");
		fclose(in);
		exit(1);
	}
	out_genes=fopen("out_cross_correlation_gene_name_table.txt","w"
);
```

```
	if(out_genes==NULL){
		printf("[ERROR] Could not open output file 
out_cross_correlation_gene_name_table.txt. Terminating.\n");
		fclose(in);
		fclose(out);
		exit(1);
	}

	//Figure out size of file and if it can be loaded into memory
	while((c=getc(in))!=13 && c!=EOF); // go to end of header line 
containing cell names
	cell_num=0;
	fscanf(in,"%*s"); //first gene name
	c=getc(in);
	while(c!=13 && c!= EOF){
		cell_num += 1;
		fscanf(in,"%*f");
		c=getc(in);
	}
	printf("[INFO] Data matrix contains %i cells.\n", cell_num);

	gene_num=1;
	while(fscanf(in,"%*s")!=EOF){
		gene_num += 1;
		while((c=getc(in))!=13 && c!=EOF); //go to end of line
	}
	printf("[INFO] Data matrix contains %i genes.\n", gene_num);

	rewind(in);

	// Load data into memory
	i = (gene_num * sizeof(char) * 40 + 
gene_num*cell_num*sizeof(float))/1024/1024;
	printf("[USER INPUT] Loading data into memory will require %i 
MB of memory.\n", i);
	printf("[USER INPUT] Proceed (y/n)? ");
	scanf("%c",&c);
	if(c!='y'){
		printf("Terminating upon user request.\n");
		fclose(in);
		fclose(out);
		fclose(out_genes);
		exit(1);
	}

	gene_names=malloc(sizeof(char)*40*gene_num);
	if(gene_names==NULL){
		printf("[ERROR] Could not allocate memory for 
gene_names.\n Terminating.\n");
		fclose(in);
		fclose(out);
		fclose(out_genes);
		exit(1);
	}
```

```
	reads=malloc(sizeof(float)*gene_num*cell_num);
	if(reads==NULL){
		printf("[ERROR] Could not allocate memory for reads.\n
Terminating.\n");
		fclose(in);
		fclose(out);
		fclose(out_genes);
		exit(1);
	}
	while((c=getc(in))!=13 && c!=EOF); // go to end of header line
containing cell names
	i=0;
	while(fscanf(in,"%39s",&gene_names[40*i])!=EOF){
		for(j=0; j<cell_num; j++){
			fscanf(in,"%f",&reads[cell_num*i+j]);
		}
		i+=1;
	}

	//Load mapping genes into memory
	mapping_genes_num=0;
	while(fscanf(in_mapping_genes,"%*i\t%*s")!=EOF){
		mapping_genes_num += 1;
	}
	rewind(in_mapping_genes);
	printf("[INFO] %i mapping genes found.\n", mapping_genes_num);
	mapping_genes=malloc(mapping_genes_num*sizeof(struct
correspondance));
	for(i=0; i<mapping_genes_num; i++){
		fscanf(in_mapping_genes,"%i\t%s", &mapping_genes[i].num,
mapping_genes[i].name);
		found = 0;
		j=0;
		while(!found && j<gene_num){

	if(!strcmp(mapping_genes[i].name,&gene_names[40*j])){
				mapping_genes[i].num_in_seq_database = j;
				found = 1;
			}
			j += 1;
		}
		if(!found){
			printf("[ERROR] mapping gene %s not found in
sequencing database !\n Terminating!\n", mapping_genes[i].name);
			exit(1);
		}
	}

	//Calculate and output results
	for(i=0; i<gene_num; i++){
		fprintf(out_genes,"%s\t%i\n", &gene_names[40*i],i);
	}
```

```
	printf("[INFO] Computing cross-correlations for genes...\n");
	for(i=0; i<gene_num; i++){
		for(l=0; l<mapping_genes_num; l++){
			j=mapping_genes[l].num_in_seq_database;
			//calculate means for genes i and j
			mean_i=0;
			mean_j=0;
			for(k=0; k<cell_num; k++){
				mean_i += reads[cell_num*i+k];
				mean_j += reads[cell_num*j+k];
			}
			mean_i = mean_i/cell_num;
			mean_j = mean_j/cell_num;

			// identify outlier
			max=0;
			outlier=0;
			for(k=0; k<cell_num; k++){
				if((reads[cell_num*i+k]-
mean_i)*(reads[cell_num*j+k]-mean_j)>max){
					max=(reads[cell_num*i+k]-
mean_i)*(reads[cell_num*j+k]-mean_j);
					outlier=k;
				}
			}

			// recalculate mean without outlier
			mean_i=0;
			mean_j=0;
			for(k=0; k<cell_num; k++){
				if(k!=outlier){
					mean_i += reads[cell_num*i+k];

					mean_j += reads[cell_num*j+k];
				}
			}
			mean_i = mean_i/(cell_num-1);
			mean_j = mean_j/(cell_num-1);
			// calculate numerator
			numerator=0;
			for(k=0; k<cell_num; k++){
				if(k!=outlier) numerator = numerator +
(reads[cell_num*i+k]-mean_i)*(reads[cell_num*j+k]-mean_j);
			}

			// calculate denominator parts
			denom_i = 0;
			denom_j = 0;
			for(k=0; k<cell_num; k++){
				if(k!=outlier){
					denom_i = denom_i +
(reads[cell_num*i+k]-mean_i)*(reads[cell_num*i+k]-mean_i);
					denom_j = denom_j +
(reads[cell_num*j+k]-mean_j)*(reads[cell_num*j+k]-mean_j);
```

```
                        }
                }

                // calculate correl coefficient
                if(denom_i*denom_j != 0){
                        correl = numerator / sqrtf(denom_i*denom_j);
                } else {
                        correl = 0;
                }

                //output
                fprintf(out,"%f", correl);
                if(l<(mapping_genes_num-1)){
                        fprintf(out,"\t");
                } else {
                        fprintf(out,"\n");
                }
            }
        }
        printf("[INFO] Done!\n");

        //Close and exit
        fclose(in);
        fclose(out);
        fclose(out_genes);
}
```

**Code to calculate expression maps**

```
#include <stdio.h>
#include <stdlib.h>
#include <string.h>
#include <unistd.h>
#include <math.h>

void WriteHexString(FILE *fptr,char *s);
void WriteTIFFfile(char *geneName, int width, int height, float
*pixels);

int main(int argc, char* argv[]){
        FILE *in_genelist, *in_mapping_genes, *in_map,
*in_correlations, *in_genesToMap;
        char gene_name[40];
        char s[40];
        int query_gene_index;
        int mapping_gene_num;
        int i, j, k;
        char c;
        float *correlations;
        int map_call;
```

```
	float score;
	float *map;
	int num_genes_to_map;
	int found;

	struct gene_list{
		char name[40];
		int found;
	} *genes_to_map;

	#define MAP_WIDTH 260
	#define MAP_HEIGHT 360

	//allocate memory for map
	if((map = malloc(sizeof(float)*MAP_WIDTH*MAP_HEIGHT))==NULL){
		printf("[ERROR] Could not allocate memory for map data.
Terminating.\n");
		exit(1);
	}

	//open files
	if((in_genelist=fopen("cross_correlation_gene_name_table.txt","
r"))==NULL){
		printf("[ERROR] Could not open input file
cross_correlation_gene_name_table.txt. Terminating.\n");
		exit(1);
	}
	if((in_mapping_genes=fopen("cross_correlation_mapping_genes.txt
","r"))==NULL){
		printf("[ERROR] Could not open input file
cross_correlation_mapping_genes.txt. Terminating.\n");
		fclose(in_genelist);
		exit(1);
	}
	if((in_map=fopen("combined_map.txt","r"))==NULL){
		printf("[ERROR] Could not open input file
combined_map.txt. Terminating.\n");
		exit(1);
	}
	if((in_correlations=fopen("cross_correlation.txt","r"))==NULL){
		printf("[ERROR] Could not open input file
cross_correlation.txt. Terminating.\n");
		exit(1);
	}

	//count mapping genes
	mapping_gene_num=0;
	while(fscanf(in_mapping_genes,"%*i\t%*s")!=EOF){
		mapping_gene_num += 1;
	}
	rewind(in_mapping_genes);
```

```
	printf("[INFO] Found %i mapping genes in file
mapping_genes.txt.\n",mapping_gene_num);

	//check mapping genes are in same order in two files
	fscanf(in_map,"%*s\t%*s"); //skip first two column names
	for(i=0; i<mapping_gene_num; i++){
		fscanf(in_map,"%s",s);
		fscanf(in_mapping_genes,"%*i\t%s",gene_name);
		if(strcmp(s,gene_name)){
			printf("[ERROR] List of mapping genes in
combined_map.txt does not correspond to list in cross-correlation
table. Terminating!\n");
			exit(1);
		}
	}
	c=getc(in_map);
	if(c!=10){
		printf("[ERROR] List of mapping genes in combined_map.txt
is longer than the mapping genes in
cross_correlation_mapping_genes.txt. Terminating\n");
		exit(1);
	}

	//check map fits with given width and height
	i=0;
	rewind(in_map);
	while(getc(in_map)!=10); //go to end of header line
	while(fscanf(in_map,"%*i")!=EOF){
		i+=1;
		while(getc(in_map)!=10);
	}
	if(i!=MAP_WIDTH*MAP_HEIGHT){
		printf("[ERROR] Map height & width do not fit available
data in combined_map.txt. Terminating.\n");
		exit(1);
	}

	//check user gave a valid file name with gene list
	if(argc != 2){
		printf("[ERROR] No input file name containing list of
genes to map found!\n");
		printf("USAGE: expression_map [name of file with list of
gene names to map]. Terminating.\n");
		exit(1);
	}
	if((in_genesToMap=fopen(argv[1],"r"))==NULL){
		printf("[ERROR] Could not open input file %s.
Terminating.\n", argv[1]);
		exit(1);
	}

	//input list of genes to map
	num_genes_to_map = 0;
	while(fscanf(in_genesToMap,"%39s", gene_name)!=EOF){
		num_genes_to_map += 1;
```

```
	}
	rewind(in_genesToMap);
	printf("[INFO] Found %i genes to map in input file.\n", 
num_genes_to_map);
	if((genes_to_map = malloc(sizeof(struct 
gene_list)*num_genes_to_map))==NULL){
		printf("[ERROR] Could not allocate memory for gene_list. 
Terminating.\n");
		exit(1);
	}
	i=0;
	while(fscanf(in_genesToMap,"%39s", gene_name)!=EOF){
		strcpy(genes_to_map[i].name, gene_name);
		genes_to_map[i].found = 0;
		i += 1;
	}

	//Allocate memory for correlation data
	if((correlations = 
malloc(sizeof(float)*mapping_gene_num))==NULL){
		printf("[ERROR] Could not allocate memory for correlation 
data. Terminating.\n");
		exit(1);
	}

	//Make directory to save output maps
	system("mkdir output_absolute_maps_directory");

	//Make maps
	while(fscanf(in_genelist,"%s\t%i",gene_name,&query_gene_index)!
=EOF){

		//check if it's in the list of genes to map
		found = 0;
		for(i=0; i<num_genes_to_map && found==0; i++){
			if(!strcmp(genes_to_map[i].name, gene_name)){
				found = 1;
				genes_to_map[i].found = 1;
			}
		}
		if(found == 0){
			while((c=getc(in_correlations)) != 10 && c!=EOF); 
// go to next row
		} else {
			for(i=0; i<mapping_gene_num; i++){

	fscanf(in_correlations,"%f",&correlations[i]);
			}
			while((c=getc(in_correlations)) != 10 && c!=EOF); 
// go to end of row

			//Calculate probability
			//NOTE : STILL NEED TO NORMALIZE FOR TOTAL 
EXPRESSION LEVELS !!
```

```
				rewind(in_map);
				while(getc(in_map)!=10); //go to end of header line
				for(i=0; i<MAP_HEIGHT; i++){
					for(j=0; j<MAP_WIDTH; j++){
						fscanf(in_map,"%*i"); // cell number
						fscanf(in_map,"%i",&map_call); // in
wing disc?
						if(map_call==0){
							map[MAP_WIDTH*i+j]=0;
							while(getc(in_map)!=10); // go to
end of this line
						} else {
							score=0;
							for(k=0; k<mapping_gene_num;
k++){

	fscanf(in_map,"%i",&map_call);
								if(map_call==0){ //mapping
gene not expressed in this cell
									score -=
correlations[k];
								}else{
									score +=
correlations[k];
								}
							}
							map[MAP_WIDTH*i+j]=score;
						}
					}
				}
				WriteTIFFfile(gene_name, MAP_WIDTH, MAP_HEIGHT,
map);
			}
	}

	//Check which genes were not found
	j=0;
	for(i=0; i<num_genes_to_map; i++){
		if(genes_to_map[i].found ==0){
			printf("[ERROR] Did not find gene %s!\n",
genes_to_map[i].name);
		} else {
			j+=1;
		}
	}
	printf("[INFO] Found %i of %i genes in input list to map.\n",
j, num_genes_to_map);

	//Close and exit
	free(map);
	free(correlations);
	fclose(in_genelist);
	fclose(in_mapping_genes);
```

```
	fclose(in_map);
	fclose(in_correlations);
	fclose(in_genesToMap);
}

void WriteHexString(FILE *fptr,char *s)
{
   unsigned int i,c;
   char hex[3];

   for (i=0;i<strlen(s);i+=2) {
      hex[0] = s[i];
      hex[1] = s[i+1];
      hex[2] = '\0';
      sscanf(hex,"%X",&c);
      putc(c,fptr);
   }
}

void WriteTIFFfile(char *geneName, int width, int height, float
*pixels){
	char filename[150];
	FILE *out;
	float min, max, range;
	int i, j, pix;
	int offset;

	strcpy(filename, "output_absolute_maps_directory/map_");
	strcat(filename,geneName);
	strcat(filename, ".tiff");

	if((out=fopen(filename,"w"))==NULL){
		printf("[ERROR] Could not open output file %s.
Terminating.\n", filename);
		exit(1);
	}

	// NORMALIZE VALUES TO RANGE 0-255
	min=pixels[0];
	max=pixels[0];
	for(i=0; i<width*height; i++){
		if(pixels[i]<min) min=pixels[i];
		if(pixels[i]>max) max=pixels[i];
	}

	if(min<-5) printf("[WARNING] Min pixel value for gene %s (%f)
is less than -5!\n", geneName, min);
	if(max>5) printf("[WARNING] Max pixel value for gene %s (%f) is
greater than +5!\n", geneName, max);
	min = -5;
	max = 5;

	range=max-min;
	for(i=0; i<width*height; i++){
```

```
            if(pixels[i]){ // exactly 0 means not in wing disc ->
leave unchanged
                  if(pixels[i]<-5) pixels[i]=-5; //for using an
absolute pixel range
                  if(pixels[i]>5) pixels[i]=5; //for using an
absolute pixel range
                  pixels[i] = (pixels[i]-min)/range*255;
            }
      }

/* Write the header */
   WriteHexString(out,"4d4d002a");    /* Big endian & TIFF identifier
*/
   offset = width * height * 3 + 8;
   putc((offset & 0xff000000) / 16777216,out);
   putc((offset & 0x00ff0000) / 65536,out);
   putc((offset & 0x0000ff00) / 256,out);
   putc((offset & 0x000000ff),out);

   /* Write the binary data */
   for (i=0;i<height;i++) {
      for (j=0;j<width;j++) {
            if(pixels[i*width + j]==0){
                  fputc(255,out);
                  fputc(255,out);
                  fputc(255,out);
            } else {
                  pix = pixels[i*width + j];
                  fputc(255-pix*235/255,out);
                  fputc(255-pix*245/255,out);
                  fputc(255-pix*131/255,out);
            }
      }
   }

   /* Write the footer */
   WriteHexString(out,"000e");  /* The number of directory entries
(14) */

   /* Width tag, short int */
   WriteHexString(out,"0100000300000001");
   fputc((width & 0xff00) / 256,out);    /* Image width */
   fputc((width & 0x00ff),out);
   WriteHexString(out,"0000");

   /* Height tag, short int */
   WriteHexString(out,"0101000300000001");
   fputc((height & 0xff00) / 256,out);    /* Image height */
   fputc((height & 0x00ff),out);
   WriteHexString(out,"0000");

   /* Bits per sample tag, short int */
   WriteHexString(out,"0102000300000003");
   offset = width * height * 3 + 182;
```

```
putc((offset & 0xff000000) / 16777216,out);
putc((offset & 0x00ff0000) / 65536,out);
putc((offset & 0x0000ff00) / 256,out);
putc((offset & 0x000000ff),out);

/* Compression flag, short int */
WriteHexString(out,"010300030000000100010000");

/* Photometric interpolation tag, short int */
WriteHexString(out,"010600030000000100020000");

/* Strip offset tag, long int */
WriteHexString(out,"011100040000000100000008");

/* Orientation flag, short int */
WriteHexString(out,"011200030000000100010000");

/* Sample per pixel tag, short int */
WriteHexString(out,"011500030000000100030000");

/* Rows per strip tag, short int */
WriteHexString(out,"0116000300000001");
fputc((height & 0xff00) / 256,out);
fputc((height & 0x00ff),out);
WriteHexString(out,"0000");

/* Strip byte count flag, long int */
WriteHexString(out,"0117000400000001");
offset = width * height * 3;
putc((offset & 0xff000000) / 16777216,out);
putc((offset & 0x00ff0000) / 65536,out);
putc((offset & 0x0000ff00) / 256,out);
putc((offset & 0x000000ff),out);

/* Minimum sample value flag, short int */
WriteHexString(out,"0118000300000003");
offset = width * height * 3 + 188;
putc((offset & 0xff000000) / 16777216,out);
putc((offset & 0x00ff0000) / 65536,out);
putc((offset & 0x0000ff00) / 256,out);
putc((offset & 0x000000ff),out);

/* Maximum sample value tag, short int */
WriteHexString(out,"0119000300000003");
offset = width * height * 3 + 194;
putc((offset & 0xff000000) / 16777216,out);
putc((offset & 0x00ff0000) / 65536,out);
putc((offset & 0x0000ff00) / 256,out);
putc((offset & 0x000000ff),out);

/* Planar configuration tag, short int */
WriteHexString(out,"011c00030000000100010000");

/* Sample format tag, short int */
WriteHexString(out,"0153000300000003");
```

```
    offset = width * height * 3 + 200;
    putc((offset & 0xff000000) / 16777216,out);
    putc((offset & 0x00ff0000) / 65536,out);
    putc((offset & 0x0000ff00) / 256,out);
    putc((offset & 0x000000ff),out);

    /* End of the directory entry */
    WriteHexString(out,"00000000");

    /* Bits for each colour channel */
    WriteHexString(out,"000800080008");

    /* Minimum value for each component */
    WriteHexString(out,"000000000000");

    /* Maximum value per channel */
    WriteHexString(out,"00ff00ff00ff");

    /* Samples per pixel for each channel */
    WriteHexString(out,"000100010001");

       fclose(out);
       return;
}
```
